# Why Meaning Survives Noise: The Spatiotemporal Abstraction Theory

**DOI:** 10.1101/2025.06.28.662092

**Authors:** Gene Fridman

**Affiliations:** Department of Otolaryngology, Johns Hopkins University, Baltimore, MD, USA; Department of Biomedical Engineering, Johns Hopkins University, Baltimore, MD, USA

## Abstract

The brain excels at extracting meaning from noisy and degraded input, yet the computational principles that underlie this robustness remain unclear. We propose a theory of spatiotemporal abstraction (STA), in which neural networks integrate inputs across space and time to produce multi-scale, concept-level representations that remain stable despite loss of detail. We demonstrate this principle using spectrograms of spoken sentences and their degraded analogs from cochlear implants, showing that as integration kernels widen, distorted input converges toward the original representation. This mechanism may explain how cochlear implant users comprehend speech despite severely scrambled afferent signals. STA provides a unified framework for understanding abstraction as an emergent property of cortical architecture, with implications for memory, neuroprosthesis design, and robust artificial systems.

---

Cochlear implants (CIs) have enabled profoundly deaf individuals to recognize speech since their introduction in the early 1970s. At the time, the concept was controversial: the electrical stimulation delivered by these devices would grossly distort the carefully orchestrated neural activity found in normal hearing. How such distorted input could be interpreted by cortical circuits adapted to clean, structured sensory input remained an open question. Yet despite this mismatch, CI users can understand sentences with remarkable success even though the neural signals reaching their auditory cortex are sparse, noisy, and highly degraded ^1,2^.

This paradox highlights a broader principle: the brain is remarkably adept at extracting meaning from incomplete, ambiguous, or distorted input. This capacity for abstraction underlies not only speech perception, but also object recognition, memory, language, and decision making ^3,4^. While hierarchical organization and functional invariance have been extensively documented across sensory and association cortices, the mechanisms by which abstraction emerges from neural architecture remain poorly understood ^5,6^.

Here we propose a theory of **spatiotemporal abstraction (STA)**, in which conceptual meaning arises through systematic integration of neural activity across space and time. In this framework, concepts are encoded as broad, distributed patterns of activity, while fine-scale details encode specific instances. The degree of abstraction depends on the scale of integration: as spatial and temporal kernels widen, representations lose detail but gain invariance.

To illustrate this principle, we examine spectrograms of spoken sentences subjected to progressive spatial blurring and compare them to quantized spectrograms that simulate CI processing. As blur increases, the spectrograms increasingly resemble their quantized counterparts—mirroring how implant users comprehend speech despite massive information loss. This convergence suggests that abstraction is not a high-level symbolic operation, but a computational consequence of hierarchical spatiotemporal integration in neural systems.

This observation motivates a general theory of how the brain encodes meaning across sensory and cognitive domains. We formalize this idea through a model of spatiotemporal abstraction, in which neural representations are shaped by the scale of integration across space and time. As integration widens, input patterns are blurred, specific details are lost, and more generalized, concept-level structure emerges. We develop this framework mathematically and demonstrate its implications using speech spectrograms, showing how coarse integration leads to convergence with cochlear implant representations. We further propose that this principle of emergent abstraction through hierarchical integration applies broadly across sensory modalities and brain regions, offering a unified account of how meaning survives noise.

## Spatiotemporal Abstraction (STA) Model

We model a cortical processing module as a population of spatiotemporal integrators that vary in their integration scale across both space and time. While this model generalizes across neural systems, it is most intuitively illustrated by the retina, where many of its core features are anatomically and functionally explicit.

Retinal ganglion cells (RGCs) form mosaics of receptive fields that vary in size and temporal dynamics, effectively tiling the visual field with overlapping layers of integration. Each RGC computes a spatiotemporal average of the photoreceptor input within its receptive field. Cells with small receptive fields capture fine detail; those with larger receptive fields produce blurred, more abstract representations. Crucially, multiple mosaics of differing spatial scales operate in parallel, allowing the same visual scene to be simultaneously encoded at different levels of abstraction ^7–10^.

We propose that similar multi-scale integration architectures exist throughout the cortex, where they have often gone unrecognized as mechanisms of abstraction. To formalize this idea, we define the input to a generic neural network as a function *I(s,t)*, where *s* represents spatial position or, more generally, a distribution over strongly weighted synaptic inputs, and *t* is time. For each output unit in the network, we define a spatiotemporal integration kernel:

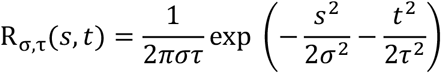

Here, *σ* and *τ* determine the spatial and temporal scales of integration, respectively. The output of a cell at a given integration scale is computed via convolution:

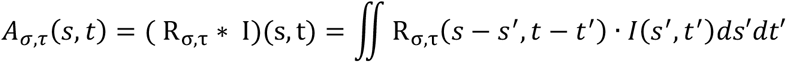

This operation filters the input through a spatiotemporal lens whose scale determines the level of abstraction. At small *σ* and *τ*, the output closely reflects local input detail; at larger scales, fine-grained features are smoothed away, leaving a more generalized encoding (**Fig. 1**). Conceptually, this defines a continuum from instance-level specificity to category-level abstraction.

**Figure 1.**
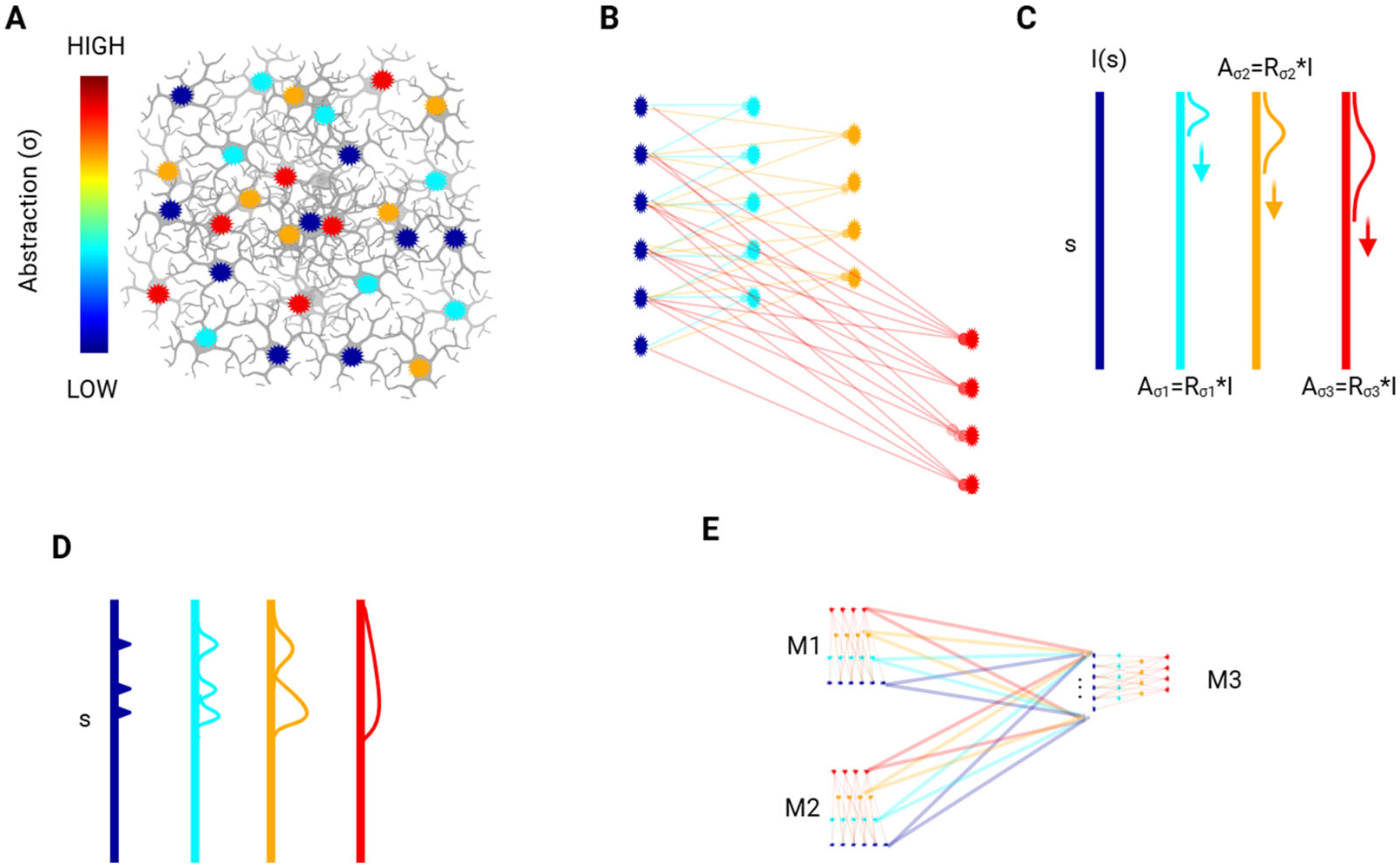
Illustration of the spatiotemporal abstraction (STA) model, shown here in its spatial form for simplicity. (**A**) Schematic of a neural module composed of interconnected units (e.g., neurons or subnetworks), color-coded by their spatial integration scale (σ). Units with larger σ integrate over broader subsets of the input population and thus encode more abstract features. (**B**) Abstracted hierarchy corresponding to panel A, where each layer represents a different spatial scale of integration applied directly to the input layer. Higher-σ units pool from more distant inputs but do not operate on each other. Red neurons are offset to illustrate the integration windows. (**C**) Mathematical formulation of STA: input signal I(s) is convolved with a spatial Gaussian kernel R_σ_(s), producing output A_σ_(s) at each abstraction level. (**D**) Example of abstraction via Gaussian convolution: a structured input (blue) is progressively smoothed into coarse representations at higher σ, illustrating the tradeoff between resolution and abstraction. (**E**) Conceptual diagram of cross-module abstraction: two modules (M1, M2), each containing their own abstraction hierarchies, project to a downstream module (M3). M3 integrates across abstraction levels and domains, supporting generalization and concept formation.

In **Figure 1A–C**, we illustrate this principle schematically. **Figure 1A** shows a network receiving input from the input level units in blue, with the higher-level integrators color-coded by weight according to their level of abstraction. **Figure 1B** redistributes the units to show the synaptic relationships, emphasizes the strongest connections that define a cell’s functional receptive field, while **Figure 1C** abstracts this pattern as a Gaussian kernel. As shown in **Figure 1D**, increasing the spatial and temporal integration scale yields a progressive transformation from detailed representations to increasingly abstract ones. **Figure 1E** indicates that the connections between modules allows for information from each abstraction layer to be incorporated into other modules’ inputs.

## Application to Cochlear Implant Perception

To illustrate how neural circuits can benefit from spatiotemporal abstraction, we examine a long-standing puzzle in auditory neuroscience: how cochlear implant (CI) users can understand speech despite severely distorted neural input ^1,2,11^. CIs restore hearing by electrically stimulating auditory nerve fibers at different points along the cochlea’s tonotopic axis. The stimulation is typically delivered as a sequence of pulses across electrode contacts, each representing a coarse frequency band ^1,12^.

This stimulation strategy introduces two major forms of distortion ^12,13^. First, electrical current spreads beyond the target site, activating neurons outside the intended spectral region. Second, the pulsatile stimulation disrupts the natural fine structure of the acoustic signal, generating phase-locked neural responses that replace the detailed temporal patterns found in normal hearing. Importantly, this degradation occurs at the **afferent neural representation level**, not in the sound waveform itself—posing a fundamental challenge to how previously learned cortical representations might continue to function. Yet, remarkably, many CI users are able to comprehend speech almost immediately after recovery from implantation surgery.

Consider a patient who lost hearing after acquiring normal language. Suppose they previously heard the sentence “The red fox jumped over the fence.” The auditory cortex would have received a structured spectrogram-like neural representation of this input, where neurons (plotted along the y-axis) encode frequency and time (x-axis) (**Fig. 2A, top left**). Through successive layers of STA-like processing, this input would be represented at progressively higher abstraction levels via spatiotemporal integration (**Fig. 2A, left column, lower rows**).

**Figure 2.**
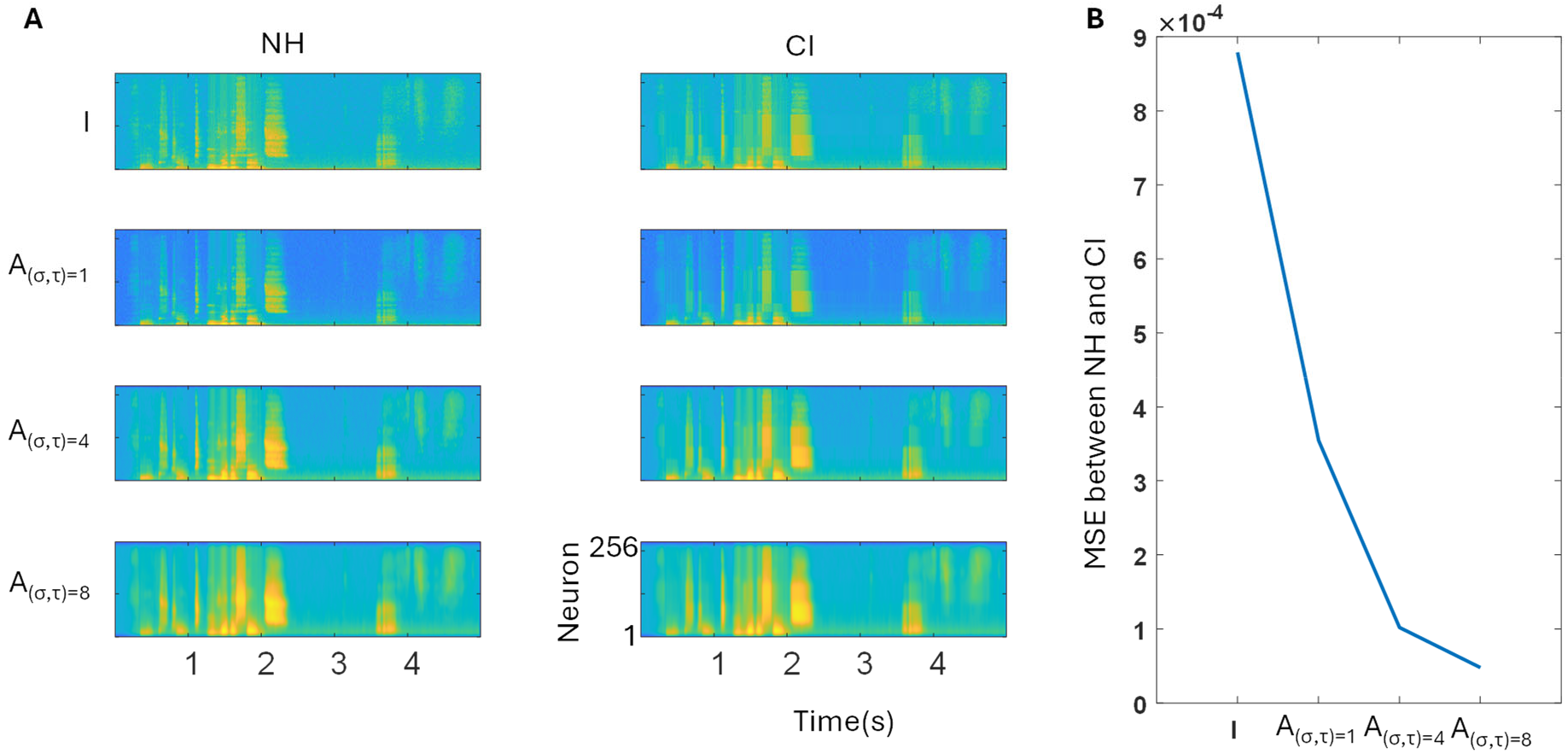
Spatiotemporal abstraction explains how cochlear implant users can recognize speech despite degraded input. (**A**) Left column: Spectrograms of a spoken sentence (“The red fox jumped over the fence”) progressively blurred using 2D Gaussian kernels with increasing spatial and temporal widths (σ = τ, from 1 to 8). These simulate hierarchical spatiotemporal abstraction layers in a normal-hearing (NH) auditory system. Each spectrogram spans 0–20 kHz on the y-axis across 256 input “neurons” logarithmically spaced by frequency, with time on the x-axis. Right column: Simulated cochlear implant (CI) input to the same network. The top panel shows an 8-channel frequency-quantized version of the original sentence, mimicking CI preprocessing. Lower rows show the resulting abstracted representations after STA integration, using the same Gaussian kernels as in the NH condition. (**B**) Mean squared error (MSE) between NH and CI representations at each abstraction level. MSE decreases monotonically as σ and τ increase, indicating that high-level abstract representations of the degraded CI input converge toward those of the original sentence. This supports the hypothesis that meaning can be preserved at coarse abstraction levels even when fine-scale input structure is lost.

After cochlear implantation, the same sentence generates a sparse, quantized, and degraded input pattern (**Fig. 2A, top right**). Yet, when passed through the original STA integration layers built from normal hearing experience, the resulting abstract representations (**Fig. 2A, right column, lower rows**) closely resemble the abstracted versions of the original sentence. We quantify this similarity using mean-squared error (MSE) across abstraction layers (**Fig. 2B**), finding that the MSE decreases monotonically with increasing integration scale. This suggests that while the low-level signal is severely distorted, meaning can still be accessed at higher abstraction levels.

To probe the robustness of this effect, we disrupted the anatomical tonotopy by randomly shuffling the input neuron order (**Fig. 3A, top left**). Despite this rearrangement, the abstraction layers still preserved similarity between the NH and CI conditions (**Fig. 3C, red and blue lines**), indicating that STA does not depend on a specific spatial layout, as long as the same connections are preserved before and after distortion.

**Figure 3.**
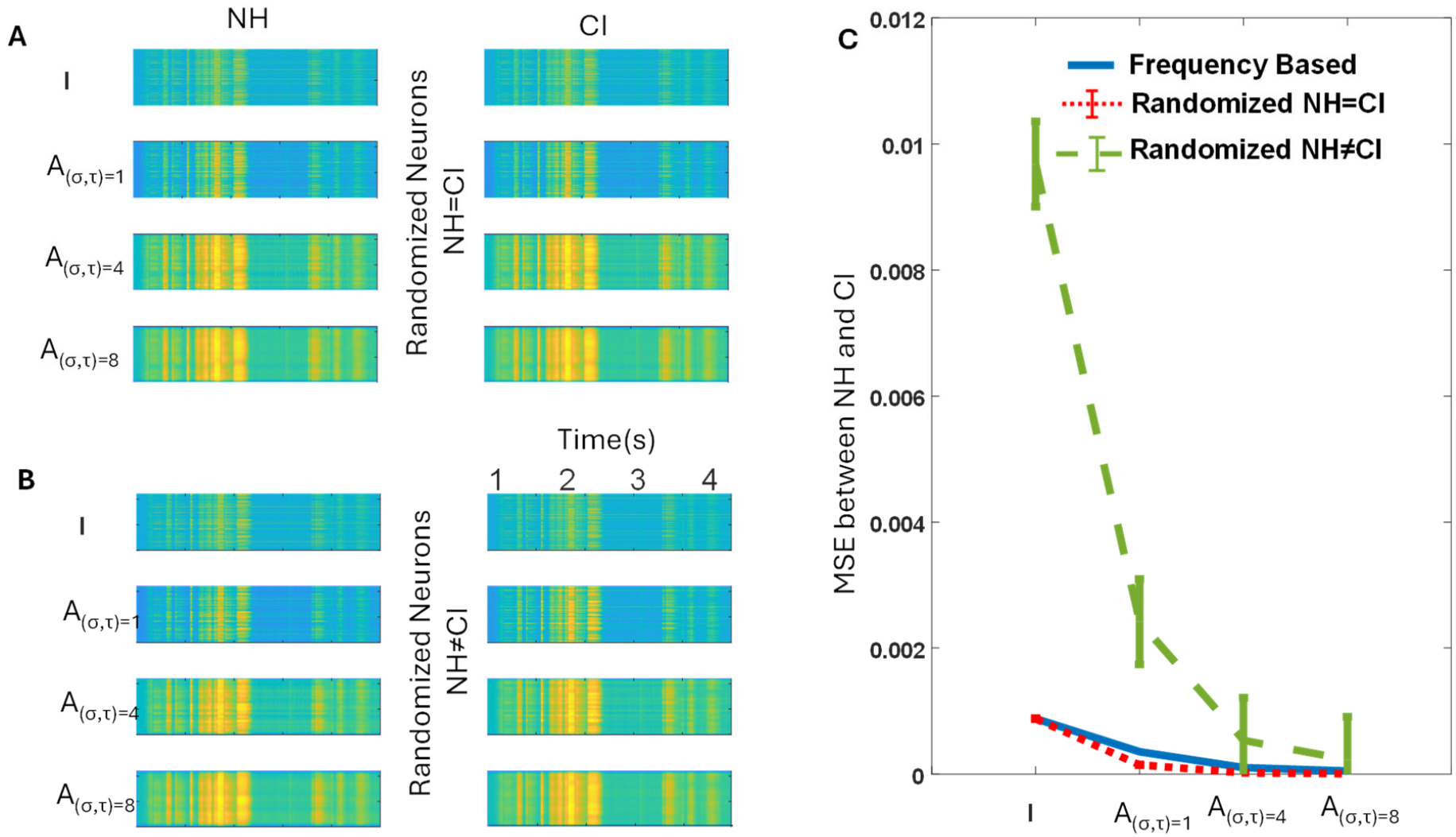
Robustness of spatiotemporal abstraction to disruption of input organization. (**A**) Input spectrograms from Figure 2 are restructured by randomly permuting the rows—removing tonotopic organization while preserving overall content. The same permutation is applied to both the normal hearing (NH) and cochlear implant (CI) conditions. Abstraction layers then operate over these shuffled input arrangements. (**B**) Control condition in which the NH and CI inputs are each randomized using different permutations, simulating a mismatch in network structure before and after implantation. This models a hypothetical scenario in which synaptic connections are reorganized following CI surgery. (**C**) Mean squared error (MSE) between the NH and CI abstraction layers at increasing blur levels (σ, τ). Blue: original tonotopic condition from Figure 2. Red: shuffled NH and CI inputs with matching permutations (from panel A, n = 10 permutations). Green: NH and CI inputs with mismatched permutations (from panel B, n = 10). Error bars denote ±1 standard deviation across permutations. Preservation of low MSE in the red condition demonstrates that STA remains effective even when spatial order is disrupted, provided the abstraction structure is consistent.

To test whether this effect depends on shared abstraction structure, we randomized the synaptic connections separately for the NH and CI inputs (**Fig. 3B**). In this case, where post-implant abstraction layers were reorganized, similarity between NH and CI representations deteriorated sharply (**Fig. 3C, green line**). However, at high abstraction levels, MSE began to decline again, approaching the original curves—suggesting that even in mismatched circuits, partial meaning may still be recoverable by chance. This may underlie the phenomenon of listeners making plausible guesses when hearing unintelligible speech.

## Biological Evidence

Spatiotemporal integration and abstraction are hallmarks of cortical processing across sensory and cognitive domains. In vision, receptive field sizes increase systematically from the retina to primary visual cortex and onward to inferotemporal areas, where neurons respond to complex objects with high tolerance to position, scale, and viewpoint changes ^3^. In audition, frequency selectivity in primary auditory cortex gives way to phonemic and linguistic encoding in higher-order areas such as the superior temporal gyrus ^14^. Similar hierarchical integration is observed in the somatosensory system ^15^, prefrontal cortex ^16^, and the entorhinal–hippocampal circuit, where place and grid cells exhibit spatial scales that vary systematically across anatomical gradients ^17^.

These findings have traditionally been interpreted as examples of abstraction, but without a unifying computational explanation. Our spatiotemporal abstraction (STA) theory proposes that abstraction naturally arises from layered integration over space and time. The retina, with its mosaic of overlapping receptive fields at multiple spatial scales, provides a concrete biological instance of this structure. While retinal ganglion cells do not directly integrate photoreceptor inputs, the interneuronal layers (bipolar, horizontal, and amacrine cells) create the necessary spatial and temporal architecture for STA-like processing.

The cochlear implant example illustrates how STA can explain a long-standing paradox: how speech recognition remains intact even when neural input is scrambled by artificial stimulation. Despite severe distortion at the input level, abstraction layers built during normal hearing can still recover meaning by integrating over time and space. The robustness of this mechanism is demonstrated by our shuffled-tonotopy simulations, where recognition performance remains intact so long as the integration structure is preserved. These findings suggest that STA confers noise invariance regardless of whether the perturbations originate internally (e.g., synaptic variability) or externally (e.g., degraded sensory input).

We suggest that the widespread anatomical and physiological evidence of spatial and temporal integration throughout the brain reflects a common computational motif. Abstraction is not a separate cognitive operation layered on top of sensory encoding; rather, it emerges directly from systematic blurring over space and time. This process supports a quantifiable tradeoff between resolution and generalization, allowing neural networks to represent both specific instances and invariant concepts. These “layers” of abstraction need not correspond to discrete cortical layers or stages but instead reflect conceptual levels of integration instantiated by synaptic weights and recurrent connectivity patterns distributed across the network.

## Broader Implications and Hypotheses

The spatiotemporal abstraction (STA) model provides a unified computational framework for understanding how the brain preserves meaning despite degraded, variable, or incomplete input. By integrating neural activity across space and time, the brain effectively smooths fine-grained variability while retaining the structural regularities that define a concept. This layered integration enables stable, high-level representations to emerge even when low-level signals are noisy or corrupted—a principle that explains, for instance, how cochlear implant users comprehend speech despite impoverished spectral input.

Beyond sensory perception, the same mechanism may underlie memory encoding. By storing abstracted, multi-scale representations, cortical circuits can preserve core meaning while discarding fragile, instance-specific details. Failures in this integration process due to atypical connectivity, temporal disorganization, or imbalanced emphasis on detail, could contribute to perceptual instability or cognitive deficits, positioning STA as a candidate framework for understanding certain neurological or psychiatric conditions.

The theory also informs the design of neuroprosthetic devices. Rather than reproducing high-fidelity sensory detail, successful prosthetics may instead aim to stimulate neural circuits in ways that align with the brain’s intrinsic abstraction hierarchies, optimizing interpretation rather than precision. Similarly, STA principles may inspire machine learning systems that emulate the brain’s ability to generalize from incomplete or noisy data. By embedding multi-scale, abstraction-based processing into artificial architectures, such systems could achieve greater robustness, energy efficiency, and interpretability in real-world environments.

In sum, STA links spatiotemporal integration to conceptual resilience, offering a biologically grounded mechanism by which meaning can survive noise. It unifies diverse findings across anatomy, physiology, and computation, and points toward a principled approach for both understanding and engineering intelligent systems. Future work should test this framework across modalities and species, develop quantitative predictions for neural encoding under degradation, and explore how manipulating abstraction hierarchies may enhance neuroprosthetic function or restore perceptual stability in clinical populations.

## Notes

### Competing Interest Statement

The authors have declared no competing interest.

